# Reptarenavirus Co- and Superinfection in Cell Culture Sheds Light on the S and L Segment Accumulation in Captive Snakes

**DOI:** 10.1101/2022.10.17.512335

**Authors:** Annika Lintala, Leonora Szirovicza, Anja Kipar, Udo Hetzel, Jussi Hepojoki

## Abstract

Boid inclusion body disease (BIBD) caused by reptarenaviruses affects collections of captive constrictor snakes worldwide. The disease manifests by formation of cytoplasmic inclusion bodies (IBs) in various tissues. Curiously, a snake with BIBD most often carries more than a single pair of genetically distinct reptarenavirus S and L segments, and the tissues of an infected individual can show variation in the variety of different S and L segment species. The role of reptarenavirus coinfection in development of BIBD remains unknown, and it is unclear how the infection affects the susceptibility to reptarenavirus superinfection or to secondary infection by other agents. Because cell culture studies with mammarenaviruses have demonstrated persistently infected cultures to resist superinfection by genetically similar viruses, we hypothesized that coinfection would only occur if the infecting viruses were of two different species. To study the hypothesis, we employed boa constrictor kidney- and brain-derived cell cultures to perform a set of co- and superinfection experiments with five reptarenavirus and one hartmanivirus isolates. Analysis of viral RNA released from coinfected cells using qRT-PCR did not demonstrate evident competition between reptarenaviruses of the same or different species in the boa constrictor kidney-derived cell line. The experiments on the brain-derived cell line revealed considerable differences in the replication ability of the reptarenaviruses tested, suggesting varying tissue specificity or target cell spectra for reptarenavirus species. Finally, experiments on persistently reptarenavirus infected cell lines showed reduced replication of closely related reptarenaviruses while the replication of reptarenaviruses from different species appeared unaffected.

**IMPORTANCE:** Snakes with boid inclusion body disease (BIBD) often display reptarenavirus coinfections, or presence of unbalanced S and L segment ratios. Studies on mammarenaviruses suggest replication interference between closely but not between more distantly related viruses. In the study, we provide evidence that similar interference or competition between segments occurs also in the case of reptarenaviruses. Conversely, the results show that there is very little or no competition between more distantly related L or S segments, the cells release similar amounts of viral RNA segments in the case of mono and coinfection. Successful superinfections of persistently infected cell cultures suggest that the unbalanced S and L segment pools often seen in the infected animals could be a result of segment accumulation through sequential reptarenavirus co- and superinfections during breeding in captivity.

## INTRODUCTION

Viruses by far outnumber their hosts (1) due to which it is not surprising that many species are coinfected by several viruses. In general, the outcomes of coinfections vary; the coinfecting viruses may e.g. compete for the same resources, in which case the dominant virus would suppress the replication of the other (2). Alternatively, coinfection is beneficial, resulting in increased replication of one or both viruses (2). The third option is that the coinfecting viruses co-exist without interfering or affecting each other’s life cycle (2). Coinfections by various pathogens (not only viruses) can have negative impact on the host’s health by amplifying the pathogenesis (3). Coinfection by closely related viruses may also lead to recombination and re-assortation of their genomes, occasionally resulting in the emergence of more pathogenic viruses (2). There are also several examples of superinfection exclusion, i.e. an established virus infection interferes or inhibits the infection by a closely related virus (2).

The family *Arenaviridae* consists of four genera, *Mammarenavirus, Reptarenavirus, Hartmanivirus*, and *Antennavirus* (4). The members of the genus *Mammarenavirus*, initially described as early as the 1930s, include mammalian arenaviruses with rodents as their natural hosts (5), with the exception of Tacaribe virus that is found in bats and ticks (6, 7). The members of the genera *Reptarenavirus* and *Hartmanivirus* infect snakes, and those of the genus *Antennavirus* infect fish (8). Arenaviruses are enveloped and have an RNA genome that is either bisegmented or, in the case of antennaviruses, trisegmented (8). The small (S) segment of the bisegmented arenaviruses encodes the glycoprotein precursor (GPC) and nucleoprotein (NP) while the large (L) segment encodes the RNA-dependent RNA polymerase (RdRp) and the matrix/Z protein (ZP) (8). Hartmaniviruses lack the ZP gene (9) as might also be the case for antennaviruses (8).

Mammarenaviruses establish an apparently benign persistent infection in their reservoir hosts (10) and can be transmitted to humans via inhalation of aerosolized rodent excreta (11). When transmitted to humans, mammarenaviruses can cause severe, even fatal disease (11) whereas the members of other arenavirus genera have not been associated to human disease. Reptarenaviruses cause boid inclusion body disease (BIBD) (12–16), a disease described in private and zoological collections of captive boid snakes since the 1970s (17) and recently also in free-ranging Costa Rican boa constrictors (18). The formation of eosinophilic intracytoplasmic inclusion bodies (IBs) in different cell types of the affected snakes is characteristic to BIBD (17). The IBs, formed at least to a major extent of reptarenavirus NP (13, 16), are more commonly seen in boas than in pythons, at least in experimental infections (12, 15). Furthermore, experimental infection (12, 15), vertical transmission (19) and screening studies performed on snake collections (20–25) suggest that reptarenaviruses establish a persistent infection at least in boa constrictors. We recently demonstrated that both reptarena- and hartmaniviruses can establish a persistent infection in cultured boid kidney cells (18).

Following the identification of reptarenaviruses, sequencing studies showed that snakes with BIBD most often carry several reptarenavirus S and L segments (26, 27). Our study further identified a coinfecting hartmanivirus in snakes with BIBD (26), but later studies suggest that hartmaniviruses do not play a role in the pathogenesis of the disease (9, 20, 28). The frequent reptarenavirus coinfection in snakes with BIBD (26, 27) and the fact that reptarenavirus S and L segments can be vertically transmitted from both parents to the offspring (19), led us to here study the coinfection dynamics of reptarena- and hartmaniviruses in cultured boa constrictor cells. By sequence comparison, the reptarenavirus S and L segments carried by BIBD-positive snakes demonstrate high divergence, and by current taxonomic criteria, a snake does not carry S nor L segments classifiable into the same reptarenavirus species. Indeed, the results of our earlier study showed that two reptarenavirus species (or two S and L segment pairs) could establish a persistent infection in cultured boid kidney cells (18), suggesting that the segments of viruses classified into two different species do not prevent each other’s replication.

We set up this study to test our hypothesis that reptarenavirus coinfection in cultured boid cells can occur with genetically dissimilar S and L segments, and that the presence of two genetically similar S and L segments will negatively affect their replication. To address the potential differences between cell types, we included two cell lines originating from different boa constrictor tissues, the kidney and the brain. We hypothesize that the frequently occurring reptarenavirus coinfections in captive snakes with BIBD could be a result of superinfection of persistently reptarenavirus-infected snakes with novel S and L segments upon introduction of new snakes into a collection. To mimic such a scenario in cell culture, we tested if the persistently reptarenavirus infected cell cultures described in our previous study (18) could be superinfected with closely or distantly related reptarenaviruses, or with a hartmanivirus. For the study, we employed five reptarenavirus and one hartmanivirus isolates, and the three persistently infected boa constrictor kidney cell lines of the earlier study. We used qRT-PCR to monitor the amount of S and L segments released from the infected cells as the readout for successive replication and virus production, and immunofluorescence staining to confirm or rule out efficient infection of the cells.

## MATERIALS AND METHODS

### Cell lines and viruses

We employed the following cell lines for studying coinfection: *Boa constrictor* kidney (I/1Ki) and *B. constrictor* brain (V/4Br) described in (16, 29). For studying superinfection, we made use of the following persistently infected cell lines: PIwUHV (positive for University of Helsinki virus 1, UHV-1, and aurora borealis virus 1, ABV-1), PIwUGV-1 (positive for University of Giessen virus 1, UGV-1), and PIwSn11 (positive for UHV-2 and Haartman Institute Snake virus 1, HISV-1), all described in (18). We maintained the cell lines at 30 °C and 5% CO_2_ in Minimal Essential Medium Eagle (MEM, Sigma Aldrich) supplemented with 10% fetal bovine serum (FBS), 200 mM L-glutamine, 100 µg/ml of streptomycin, and 100 U/ml of penicillin.

We selected the following reptarenavirus isolates: UHV-1 (GenBank accession numbers KR870011 and KR870020) (26), UHV-2 (KR870016 and KR870030) (9, 26), UGV-1 (KR870012 and KR870022) (26), UGV-2 (KR870015 and KR870029) (26)), ABV-1 (KR870010 and KR870021), and HISV-1 (KR870017 and KR870031) (9) for the study. After initial testing of coinfections with two different virus input amounts, 100 and 1000 S segment copies/cell (quantified as described below), we selected 100 S segment copies/cell which corresponds roughly to MOI (multiplicity of infection) 5-10, for the subsequent experiments conducted on cells grown on 6-well plates. The viruses diluted in fully supplemented growth medium were allowed to adsorb for an hour at 30 °C with 5% CO_2_ while gently tilting the plates every 15 minutes, after which the cells were washed three times with fully supplemented medium, fresh medium added, and the plates incubated at 30 °C with 5% CO_2_ up to eight days. Samples from the medium were collected at 0, 4 and 8 days post infection (dpi) and stored at −80 °C.

### RNA extraction and quantitative reverse transcription-polymerase chain reaction (qRT-PCR)

The GeneJET RNA purification kit (ThermoFisher Scientific) used according to the manufacturer’s protocol served for extracting RNA from cell culture supernatants. Synthetic genes under T7 promoter (Gene Universal) served for generation of control RNAs as described (18). The *in vitro* transcribed control RNAs served for translating cycle threshold (Ct) values to copy numbers via http://endmemo.com/bio/dnacopynum.php, in short, each qRT-PCR run included a 10-fold dilution series (e.g. 100,000,000 to 100,000 copies/reaction) of the respective control RNAs in duplicate. The primers and probes as well as the qRT-PCR protocol for quantifying the S and L segments and housekeeping gene were as described (18). In brief, the reactions with Taqman Fast Virus 1-step master mix (ThermoFisher scientific) were half of the recommended 20 μl volume, and the reaction components were: 2.5 μl of the master mix, 2.5 μl of template, 0.25 μl of each primer (20 μM stock), 0.125 μl of the probe (20 μM stock), and 4.375 μl of water. The cycling (1: 5 min at 50°C; 2: 20 sec at 95°C; 3: 3 sec at 95°C; 4: 30 sec at 60°C (steps 3 and 4 were repeated 40 times) was done with the AriaMX real-time PCR System (Agilent).

### Immunofluorescence (IF) staining

To study the ability of the different isolates to infect and spread in the two selected cell lines, the I/1Ki and V/4Br cells were plated on sterile ViewPlate-96 black Tissue culture plates (Perkin Elmer) coated overnight at 4 °C with 0.1 mg/ml Collagen Type 1 from rat tail (BD Biosciences) in 0.25 mM acetic acid. After overnight attachment, the cells were inoculated with the different reptarenavirus and hartmanivirus isolates as described above. At 2 or 5 dpi the growth medium was removed and the cells were fixed with 4% paraformaldehyde (PFA) in phosphate-buffered saline (PBS) for 15 min at room temperature (RT). After fixation, the cells were washed once with PBS, blocked and permeabilized for 5 min with 3% bovine serum albumin (BSA), 0.25% Triton-X-100 in Tris-buffered saline (TBS), and washed once with TBS. The cells were incubated with primary antibodies [rabbit anti-UHV NP-C or rabbit anti-HISV NP antisera, described in (9, 26) diluted 1:2,000 in TBS supplemented with 0.5% BSA for 1.5 h at RT, washed three times with TBS, incubated with secondary antibodies [Alexa Fluor 594- or 488 labelled goat anti-rabbit or donkey anti-rabbit (ThermoFisher Scientific)] diluted 1:1,000 in TBS supplemented with 0.5% BSA and 0.5 μg/ml of Hoechst 33342 for 45 min at RT, and washed 3-5 times with TBS prior to imaging. The staining was recorded either using the ZOE Fluorescent Cell imager (Bio-Rad) or the Opera Phenix High Content Screening System (PerkinElmer).

## RESULTS

### The ratio of S and L segments released into the growth medium varies between virus stocks

In our previous study, using the virus segment-specific one-step qRT-PCR, we observed varying S and L segment ratios within the cells infected with different reptarenavirus isolates (18). Therefore, we first analyzed the S and L segment RNA levels in the virus stocks utilized in this study (Table 1). The S segment RNA levels of the stocks varied from 5.5 * 10^6^ to 4.9 * 10^7^ copies per ml, and for most of the reptarenavirus (and the hartmanivirus included) isolates the S to L segment ratio remained between 1.0 and 2.8. Curiously, the University of Giessen virus 2 (UGV-2) and aurora borealis virus 1 (ABV-1) stock showed drastically higher S to L segment ratios of 6.1 and 28.6 (Table 1). Although RT-PCR efficiency could contribute to the observation, the results of our earlier study showed S to L segment ratio of 0.9 to 1.9 for ABV-1 within the cells, thus suggesting that the results would indeed reflect differences in release of segments from the infected cells.

**Table 1.** Copy number (CN) and the ratio of S segment to L segment in the virus stocks. The CNs were determined using qRT-PCR and are expressed by CN per µl. The S/L ratio expresses the molar excess of S segment to L segment in each stock.

| | S segment CN/$\mu\text{l}$ | L segment CN/$\mu\text{l}$ | Ratio (S/L) |
| --- | --- | --- | --- |
| **UHV-1** | $2.3 \times 10^7$ | $1.55 \times 10^7$ | 1.57 |
| **UHV-2** | $6.09 \times 10^6$ | $5.99 \times 10^6$ | 1.02 |
| **UGV-1** | $4.91 \times 10^7$ | $1.76 \times 10^7$ | 2.78 |
| **UGV-2** | $5.50 \times 10^6$ | $8.96 \times 10^5$ | 6.14 |
| **ABV-1** | $6.84 \times 10^6$ | $2.39 \times 10^5$ | 28.64 |
| **HISV-1** | $1.98 \times 10^7$ | $1.45 \times 10^7$ | 1.36 |

In the earlier study, we found that ABV-1 and University of Helsinki virus 1 (UHV-1) can coinfect boa constrictor kidney cells (I/1Ki cell line, (16)) without apparent competition between the viruses (18). To test whether the ability to establish coinfection would depend on the virus isolate, we inoculated I/1Ki cells with UHV-1 and UGV-1 alone or together. In addition, we tested whether the amount of S and L segment copies used for the inoculation would have an effect on the amount of S and L segments released. We used 100 S segment copies (roughly corresponding to a multiplicity of infection (MOI) of 5-10) for both UHV-1 and UGV-1 in monoinfections, and 100 or 1,000 UHV-1 S segment copies in coinfections. The results show that coinfection did not markedly alter the amount of virus RNA released (Fig 1), suggesting very little or no competition between the viruses. While using 10x more UHV-1 S segment RNA in the inoculum did not markedly affect the release of UHV-1 L segment RNA, the “imbalanced” coinfection resulted in the release of a slightly higher amount of UHV-1 S segment RNA and a slightly lower amount of UGV-1 segment RNA (1:3.5x for S and 1:2.3 for L) (Fig 1).

**Figure 1.**
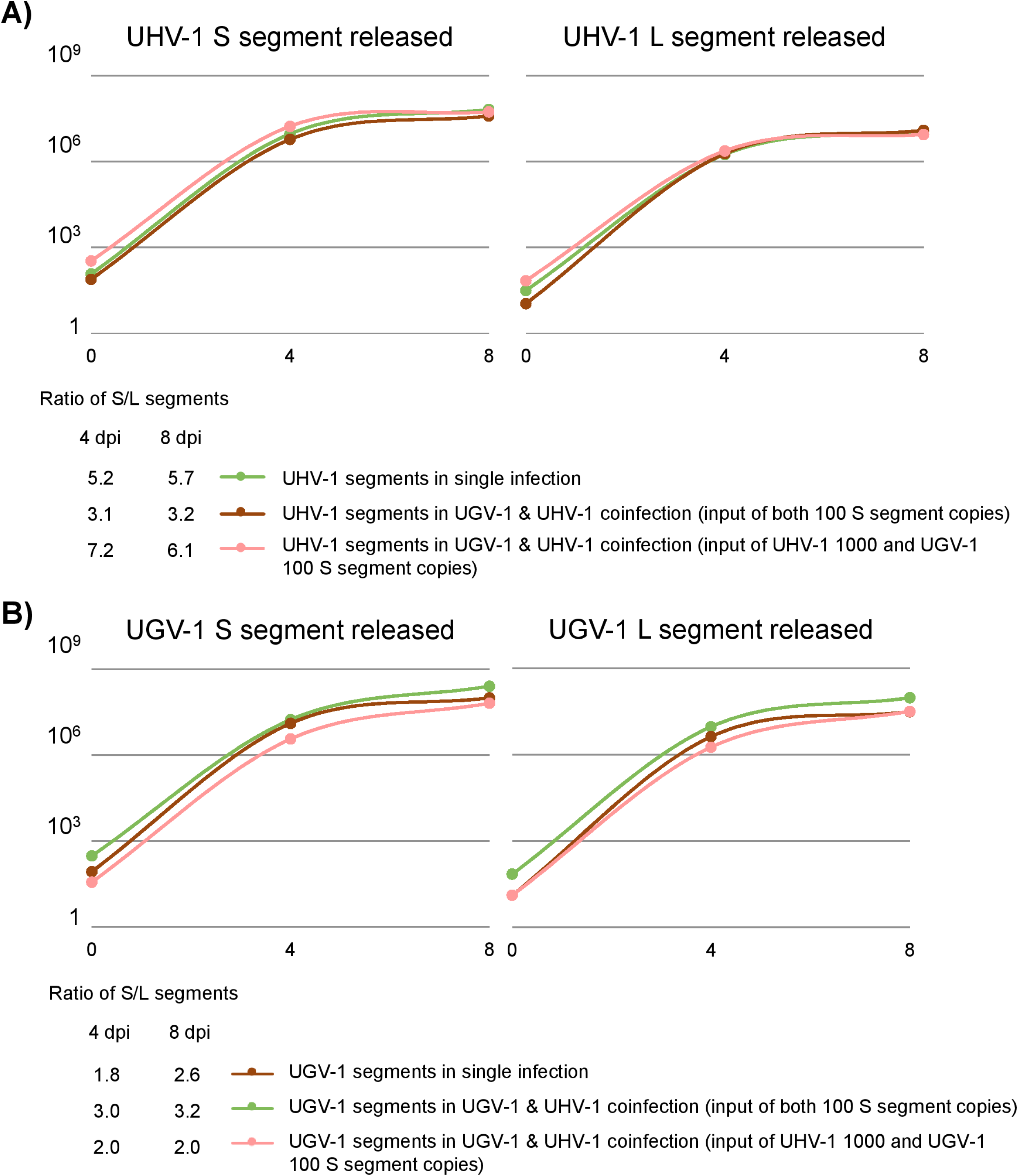
The effect of input virus on the amount of S and L segments released from the infected cells. S and L segment copy numbers (CN) in the growth medium at 4 and 8 days post inoculation (dpi) of I/1Ki cells **A)** UHV-1 S and L segments after inoculation with UHV-1 alone (100 S segment copies/cell), green line; UHV-1 and UGV-1 together at equal amounts (both 100 S segment copies/cell), dark red line; or with 10-fold excess of UHV-1 over UGV-1 (1000 S segment copies/cell of UHV-1 and 100 S segment copies/cell of UGV-1), pink line. Y-axis represents UHV-1 segment copies per 1 µl of culture medium. The table below shows the UHV-1 S/L segment ratio at each time point. **B)** UGV-1 S and L segments after inoculation with UGV-1 alone (100 S segment copies/cell), green line; UGV-1 and UHV-1 together at equal amounts (both 100 S segment copies/cell), dark red line; or with 10-fold excess of UHV-1 over UGV-1 (1000 S segment copies/cell of UHV-1 and 100 S segment copies/cell of UGV-1), pink line. Y-axis represents UGV-1 segment copies per 1 µl of culture medium. The table below shows the UGV-1 S/L segment ratio at each time point.

### Brain-derived V/4Br and kidney-derived I/1Ki boa constrictor cell lines differ in their ability to support reptarena- and hartmanivirus growth, with little or no competition during coinfection

Snakes with BIBD are commonly positive for several reptarenavirus S and L segments (19, 20, 26–28, 30), and can be in about 40-50% of cases positive for hartmaniviruses (20). The reptarenavirus S and L segments within an individual are often genetically divergent, each S and L segment representing a different reptarenavirus species. Reptarenaviruses and especially IBs can generally be detected also in the brain of boa constrictors with BIBD (13, 16, 17, 31), however, one of our earlier studies suggested that it might harbor a different set of S and L segments than the blood in an individual infected animal (19). Still, another earlier study from our group employing vesicular stomatitis virus (VSV) pseudotyped with various reptenarenavirus glycoproteins found our brain-derived boa constrictor cell line, V/4Br, to be reptarenavirus permissive (32). So far, our boa constrictor kidney-derived I/1Ki cell line seems to efficiently support the growth of all reptarenaviruses and hartmaniviruses as judged by virus isolation attempts (9, 15, 16, 19, 26, 29, 30). Therefore, to study the ability of reptarenaviruses to replicate in cells originating from different *Boa constrictor* tissues and the potential effects of coinfection, we inoculated V/4Br and I/1Ki cells with combinations of different reptarenaviruses (ABV-1, UHV-1, UHV-2, UGV-1, and UGV-2) and a hartmanivirus (Haartman Institute snake virus 1, HISV-1). We used qRT-PCR to monitor the amount of S and L segments in the cell culture supernatants collected at 4 and 8 days post inoculation (dpi) with a single virus or different pairs of two viruses. We hypothesized that in coinfection two closely related reptarenaviruses would compete in replication with each other while more distantly related reptarenaviruses would not.

The results (Fig 2) indicate that for all viruses studied, the V/4Br cells release lower amount of viral RNA into the cell culture supernatant as compared to I/1Ki cells. Depending on the virus isolate, I/1Ki cells released approximately 10-to 10,000-fold more viral RNA than V/4Br cells. Of the reptarenaviruses, UHV-1 and UHV-2 appeared to replicate the most efficiently in the V/4Br cells. Curiously, HISV-1 did not appear to replicate efficiently in V/4Br cells (Fig 2F) although our earlier study detected hartmaniviruses antigen in neurons in the brain of naturally infected boas, suggesting that it replicates in neurons *in vivo* (9). Continuing the infection up to 22 dpi did not markedly increase the amount of HISV-1 released into the medium. The results further show that coinfection did not have a clear effect to the amount of S and L segment RNA released from the infected cells into the growth medium for the majority of the tested virus combinations in I/1Ki cells. Coinfection of I/1Ki with UHV-1 and UHV-2 resulted in a 10-fold decrease in the release of UHV-1 segments as compared to single virus infection (Fig 2A). Similarly, the coinfection of UHV-1 with UGV-1 reduced the amount of UHV-1 segments released, albeit less substantially than with UHV-2. However, none of the coinfecting viruses appeared to affect the amount of UHV-2 segment released (Fig 2B). Coinfection with UGV-1 and UHV-2 caused a less than 10-fold reduction in the amount of UGV-1 segments released as compared to monoinfection (Fig 2C). The slight variation observed for segments released in the 8 dpi samples of most I/1Ki coinfection combinations most likely reflects a method-induced variation rather than true differences, suggesting that coinfection does not markedly affect reptarena- and/or hartmanivirus replication in this cell line. However, in the V/4Br cells coinfection had a larger impact on the amount of segments released for some of the tested viruses. For instance, less UHV-1 segments were released into the growth medium when in coinfection with UHV-2, ABV-1, UGV-1 or UGV-2 (Fig 2A). Supporting the initial hypothesis, coinfection with UHV-2 had the highest impact on the segment release as compared to UHV-1 monoinfection. As observed with I/1Ki cell, the hartmanivirus (HISV-1) coinfections did not affect the reptarenavirus segment release in V/4Br cells (Fig 2F).

**Figure 2.**
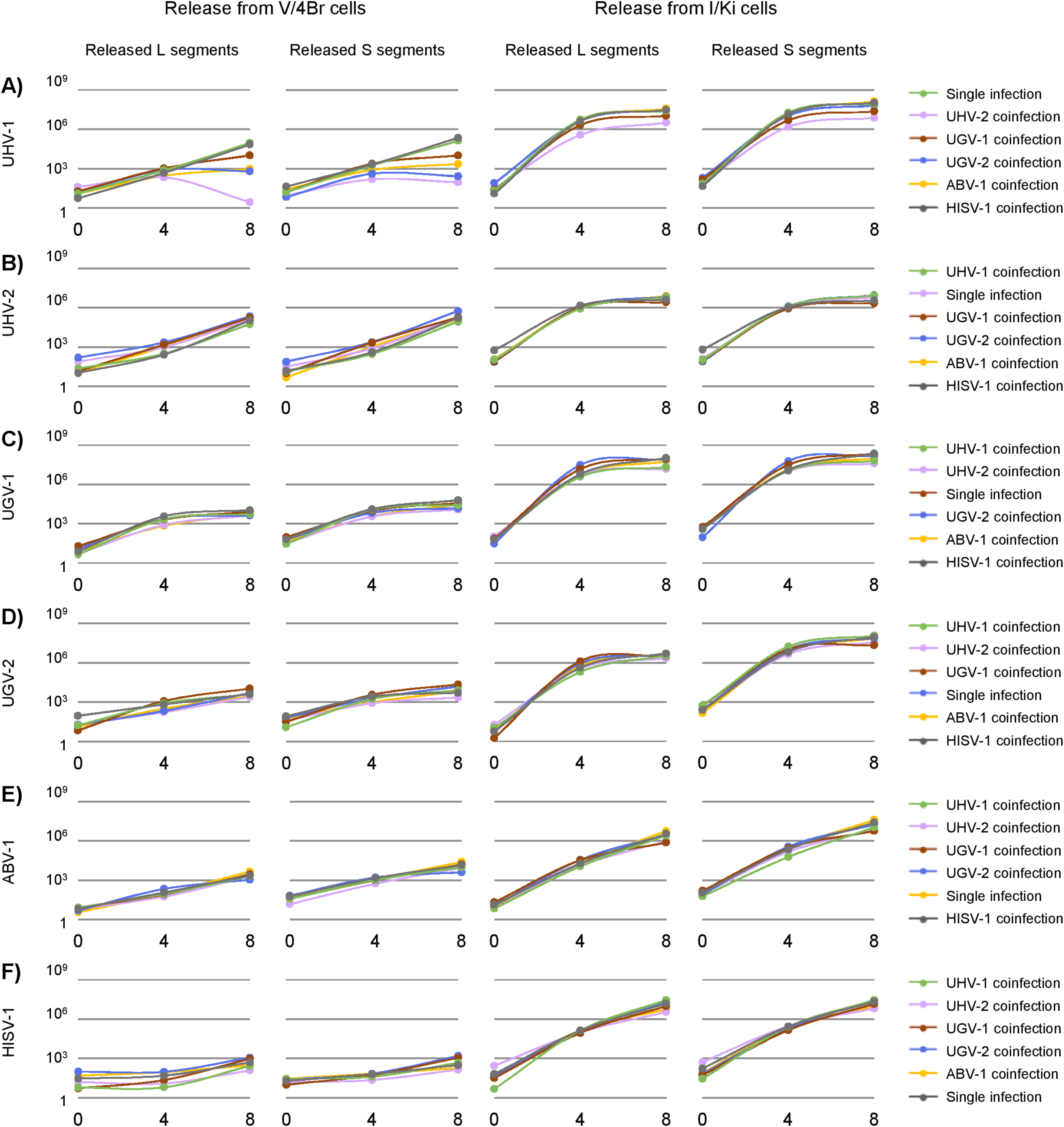
The effect of coinfection on the release of reptarena- and hartmanivirus S and L segments from V/4Br and I/1Ki cells. S and L segment RNA copy numbers (CN) in the growth medium of V/4Br (*B. constrictor* brain-derived, left panels) and I/1Ki (*B. constrictor* kidney-derived, right panels) cells inoculated either with a single virus or with pairs of two viruses. The Y-axis represents the CN of genome segments in 1 µl of the growth medium collected at 4 or 8 dpi. The first two panels on the left represent the segments released from V/4Br cells, the two panels on the right represent the segments released from I/1Ki cells, the legends on the right indicate the color coding of the lines in the graphs. CN in single virus infection versus coinfections with the indicated virus pairs are shown for the L and S segments of **A)** UHV-1, **B)** UHV-2, **C)** UGV-1, **D)** UGV-2, **E)** ABV-1, and **F)** HISV-1.

To study if the observed differences between the I/1Ki and V/4Br cell lines could be explained by differences in the ability of the different viruses to infect the cells, we used immunofluorescence staining to identify the number of infected cells at 2 and 5 dpi. While all selected virus isolates, except ABV-1, appeared to cause almost 100% infection in I/1Ki cells by day 5, only UHV-2, UGV-2, and ABV-1 had efficiently infected V/4Br cells by day 5 (Fig 3). The results of the IF staining are in line with the qRT-PCR results, and indicate that the selected virus isolates show differences in their ability to infect and replicate in V/4Br cells.

**Figure 3.**
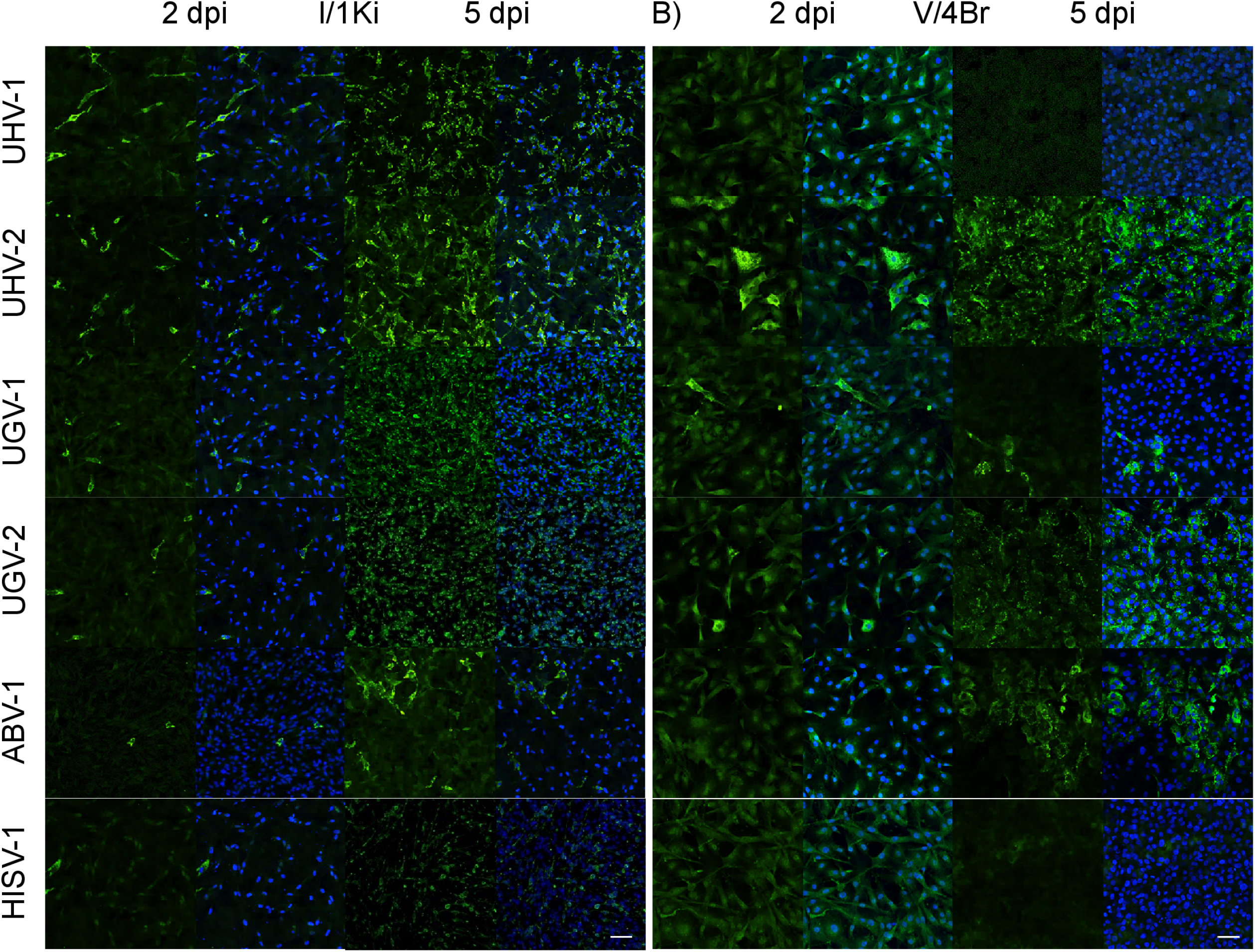
Immunofluorescence staining of I/1Ki and V/4Br cells at two and five days post inoculation with the selected viruses. I/1Ki (*B. constrictor* kidney-derived) and V/4Br (*B. constrictor* brain-derived) cells inoculated with the indicated viruses were fixed at 2 or 5 dpi and stained using antiserum against the NP (green) to detect infected cells and Hoechst 33342 to visualize the nuclei (blue): **A)** I/1Ki cells and **B)** V/4Br cells. The figures are arranged so that for each time point the green channel is shown first followed by an overlay of the green and blue channels. The images were captured with Opera Phenix, scale bar 200 µm.

### Coinfection does not affect the ratio of L and S segments released

The frequent reptarenavirus coinfections in boa constrictors with BIBD (26, 27) suggest that reptarenavirus S and L segments might re-assort rather freely (27). As studied above, the S to L segment ratio in the supernatant appeared to differ from the ratio observed in the infected cells in our earlier study (18). We thus studied whether coinfection would alter the ratio of S and L segments released. Table 2 shows the S to L segment ratio in supernatants of mono- and coinfected cells collected at 4 and 8 dpi. The results show that for some viruses, e.g. UHV-1, UGV-1, and HISV-1, the segment ratio appears to vary between the two cell lines tested. In addition, for some viruses, e.g. ABV-1 and UGV-2, the ratio appears to change markedly between the two time points studied. To better enable comparison of the segment ratio between mono- and coinfection, we calculated the average ratios from all experiments and time points for both cell lines. With some virus combinations (e.g. UHV-1 in I/1Ki cells with UGV-1 or UGV-2) a two-fold difference in the segment ratio was observed but in the majority of combinations the ratio of released S and L segments did not vary substantially between mono- and coinfection. Several technical factors can influence the results and thus we are careful with our interpretation, however, coinfection with UGV-1 in I/1Ki cells appears to affect the ratio of segments released for the majority of reptarenaviruses tested. When comparing the results to those obtained from the virus stocks in Table 1, we observed slight differences between the stocks. The virus stocks are a pool of medium collected at different time points after the infection (3-6 dpi, 6-9 dpi, and 9-12 dpi), which could explain the difference between the stock and the 4 dpi and 8 dpi samples. However, this could still indicate a change of S and L segment ratios during the course of infection, at least for some of the viruses.

**Table 2.** The ratio of S segment to L segment released from I/1Ki and V/4Br cells to the culture medium at 4 and 8 dpi with either one (single virus infection) or two viruses (coinfection). The S/L ratios presented in the table are derived by dividing the measured S segment copy number (CN) in the medium with the respective L segment CN.

|  |  | Single virus |  | UHV-1 coinf. |  | UHV-2 coinf. |  | UGV-1 coinf. |  | UGV-2 coinf. |  | ABV-1 coinf. |  | HISV-1 coinf. |  | Average |
| --- | --- | --- | --- | --- | --- | --- | --- | --- | --- | --- | --- | --- | --- | --- | --- | --- |
|  |  | 4 dpi | 8 dpi | 4 dpi | 8 dpi | 4 dpi | 8 dpi | 4 dpi | 8 dpi | 4 dpi | 8 dpi | 4 dpi | 8 dpi | 4 dpi | 8 dpi |  |
| I/1Ki | UHV-1 | 3.9 | 4.1 | - | - | 4.2 | 2.8 | 2.4 | 2.0 | 2.4 | 2.2 | 3.7 | 5.3 | 3.4 | 3.7 | 3.3 |
|  | UHV-2 | 1.3 | 1.3 | 1.5 | 1.6 | - | - | 0.7 | 0.8 | 1.1 | 1.3 | 0.9 | 1.0 | 0.8 | 1.1 | 1.1 |
|  | UGV-1 | 2.3 | 2.6 | 3.5 | 5.3 | 2.3 | 2.4 | - | - | 2.0 | 1.8 | 2.4 | 1.7 | 2.9 | 3.6 | 2.7 |
|  | UGV-2 | 4.9 | 4.0 | 7.6 | 4.6 | 3.8 | 2.9 | 4.0 | 2.1 | - | - | 4.0 | 3.9 | 4.9 | 4.1 | 4.2 |
|  | ABV-1 | 14.0 | 7.2 | 9.4 | 8.3 | 11.7 | 7.0 | 10.0 | 8.0 | 10.0 | 6.0 | - | - | 12.1 | 7.1 | 9.2 |
|  | HISV-1 | 2.5 | 2.0 | 2.0 | 1.4 | 3.0 | 2.1 | 3.8 | 2.4 | 1.8 | 1.3 | 3.1 | 2.7 | - | - | 2.3 |
| V/4Br | UHV-1 | 2.5 | 1.5 | - | - | 0.7 | 32.8 | 2.3 | 1.0 | 0.5 | 0.4 | 2.8 | 2.4 | 4.6 | 3.3 | 4.6 |
|  | UHV-2 | 1.3 | 1.3 | 1.0 | 1.5 | - | - | 1.4 | 1.1 | 1.1 | 2.5 | 1.0 | 1.1 | 1.5 | 1.6 | 1.4 |
|  | UGV-1 | 4.8 | 4.6 | 5.2 | 4.8 | 4.5 | 2.8 | - | - | 2.9 | 3.6 | 5.5 | 5.2 | 3.5 | 5.7 | 4.4 |
|  | UGV-2 | 8.9 | 3.1 | 2.1 | 2.1 | 4.8 | 0.9 | 3.0 | 2.0 | - | - | 3.1 | 2.4 | 4.0 | 1.2 | 3.1 |
|  | ABV-1 | 12.2 | 5.4 | 11.5 | 4.8 | 9.7 | 4.3 | 9.7 | 4.3 | 7.4 | 3.6 | - | - | 12.8 | 6.3 | 7.7 |
|  | HISV-1 | 1.3 | 0.6 | 6.3 | 1.7 | 1.8 | 1.2 | 2.4 | 1.2 | 0.7 | 1.4 | 0.8 | 0.6 | - | - | 1.7 |

### Superinfection of persistently infected cell lines suggests competition between closely related reptarenaviruses

We reported earlier that reptarenaviruses are transmitted vertically, i.e. when boa constrictors with BIBD reproduce; the offspring can obtain S and L segments from one or both parents (19). Experimental infection studies imply that reptarenaviruses cause a persistent infection at least in boa constrictor (12, 15), and we recently demonstrated establishment of persistent reptarenavirus infection in I/1Ki cells (18). The fact that boa constrictors with BIBD usually carry S and L segments of several reptarenavirus species suggests that superinfection of persistently infected animals might be a frequent event. To study if persistently infected cell cultures are permissive to superinfection, we made use of I/1Ki cells persistently infected (PI) with UGV-1 (PIwUGV-1), ABV-1 and UHV-1 (PIwUHV), and UHV-2 and HISV-1 coinfected (PIwSn11) described in (18). We superinfected the cell lines with the virus isolates used above, excluding the viruses used to establish the persistent infection, and monitored the amount of released L and S segments. Before initiating the experimental work, we confirmed that the PI cells were infected only with the viruses used for establishing the persistent infection using qRT-PCR for the virus isolates used in this study.

First, we studied whether superinfection of the persistently infected cell lines would affect the release of the viruses used to establish the persistent infection. We employed only S segment specific qRT-PCR, since the results above had indicated that the putative changes affect the release of both segments. The results show that superinfection did not have a notable effect on the release of the persistently infecting viruses in the PIwSn11 and PIwUHV cell lines (Fig 4A-B) coinfected with two viruses. However, superinfection of the PIwUGV-1 cells persistently infected with a single reptarenavirus (UGV-1) with all other viruses tested resulted in a >10-fold increase in the release of the persistently infecting UGV-1 (Fig 4C).

**Figure 4.**
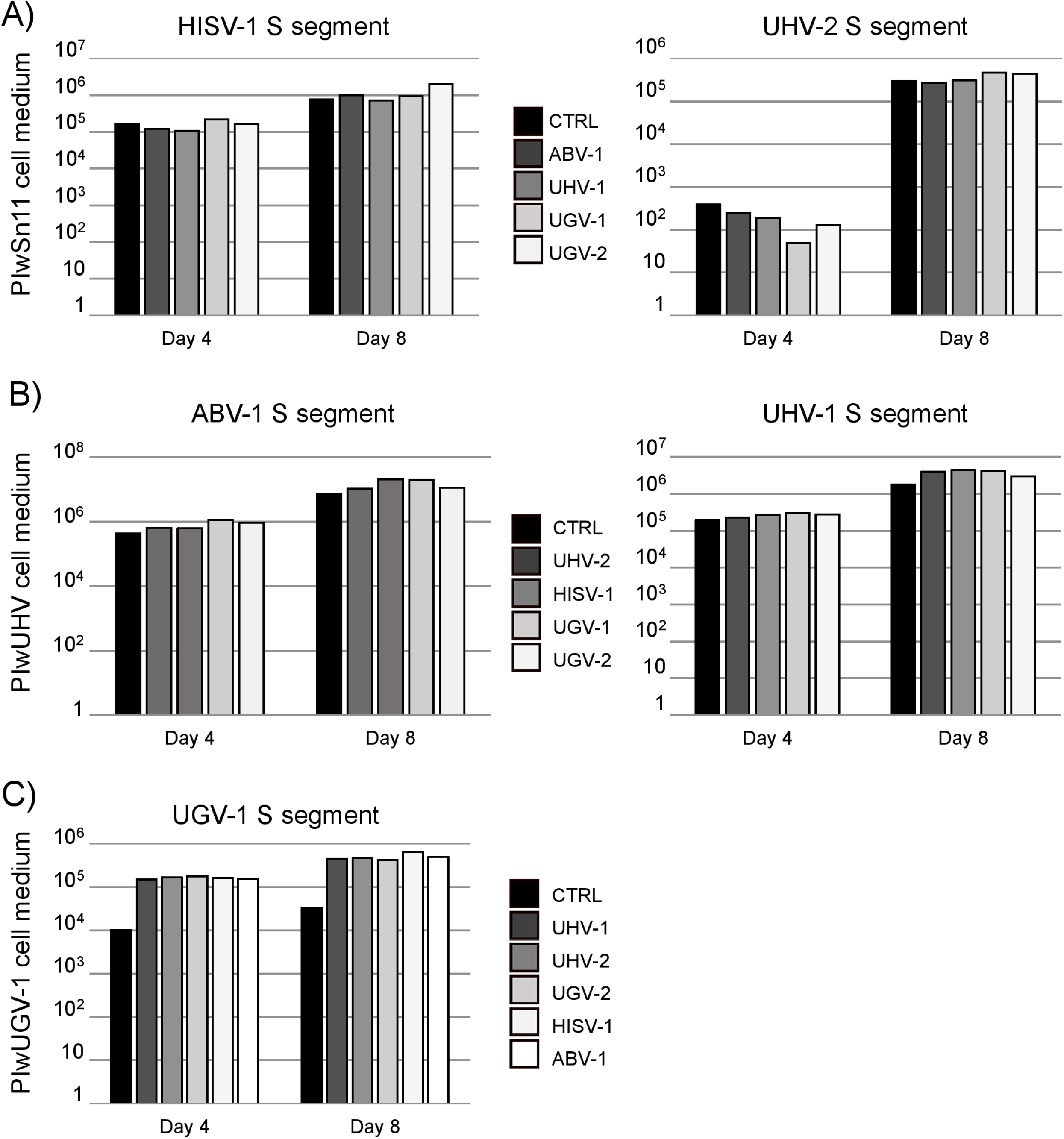
The effect of superinfection on viral RNA release in persistent infections. The supernatants of persistently infected (PI) cell lines, PIwSn11 (infected with HISV-1 and UHV-2), PIwUHV (infected with ABV-1 and UHV-1), and PIwUGV-1 (infected with UGV-1) described in (18), collected at 4 and 8 dpi following superinfection with the viruses indicated were analyzed for the amount of S segment RNA of the persistently infecting virus(es). The Y-axes of the graphs indicate copy numbers (CN) of S segment RNA per microliter of growth medium. The control (CTRL) represents non-superinfected cells. **A)** The left panel shows the HISV-1 and the right panel the UHV-2 S segment RNA released from PIwSn11 cells following superinfection. The legend in the middle shows the color coding of the bars. **B)** The left panel shows the ABV-1 and the right panel the UHV-1 S segment RNA released from PIwUHV cells following superinfection. The legend in the middle shows the color coding of the bars. **C)** UGV-1 S segment RNA released from PIwUGV-1 cells following superinfection. The legend on the right shows the color coding of the bars.

Next, we studied the release of the S and L segments of the superinfecting viruses at 4 and 8 dpi. In most virus combinations, the superinfecting viruses released their segments at a level equal or higher to that of the persistently infecting virus (indicated by the arrows in Fig 5). The results show that UGV-2 could effectively complete its infectious cycle as judged by RNA release form two of the persistently infected cell lines, PIwSn11 (Fig 5A) and PIwUHV (Fig 5B), but not from the PIwUGV-1 cells (Fig 5C). Infection of the PIwUGV-1 cells with the closely related UGV-2, resulted in release of detectable amount of UGV-2 S segment at both time points, but the UGV-2 L segment was only detected in the supernatant collected at 8 dpi. The second pair with two closely related reptarenaviruses, PIwUHV cells superinfected with UHV-2, showed also lower release of the superinfecting virus as compared to monoinfection (compare Fig 5B and 5C). Similarly, the release of UHV-1 segments from the PIwSn11 (UHV-2 and HISV-1 infected) cells following superinfection appeared to be reduced, further supporting the hypothesis of competition between the closely related viruses (compare Fig 5A and 5B). A comparison with the results obtained from infections of naïve I/1Ki cells showed roughly a 10 to 100-fold lower amount of segments released for the superinfecting virus in most cases (compare Fig 5 to Fig 2). The results indicate that the competition between closely related viruses is more evident in the superinfection setting as compared to the set up when the cells are coinoculated with the two viruses.

**Figure 5.**
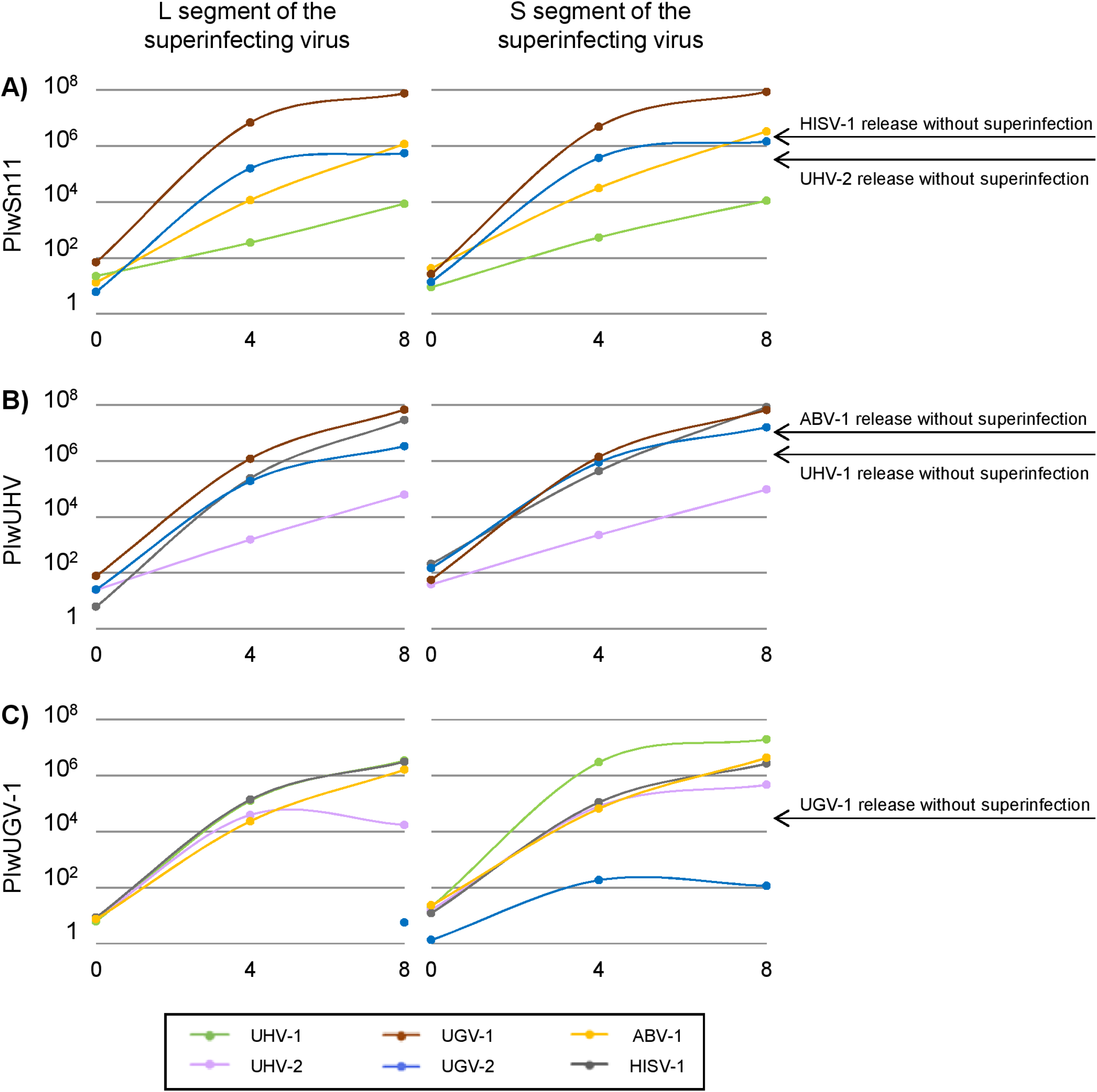
The release of S and L segment RNA following superinfection of persistently infected boid cell cultures. The supernatants of PI cell lines, PIwSn11 (infected with HISV-1 and UHV-2), PIwUHV (infected with ABV-1 and UHV-1), and PIwUGV-1 (infected with UGV-1) described in (18), collected at 4 and 8 dpi following superinfection with the viruses indicated in the panel below were analyzed for the amount of S and L segment RNA of the superinfecting infecting virus(es). The Y-axes of the graphs indicate S and L segment copy numbers (CN) per microliter of growth medium. The legend at the bottom of the figure shows the color coding of the lines. **A)** The left panel shows the amount of L and the right panel the amount of S segment RNA released from PIwSn11 cells. **B)** The left panel shows the amount of L and the right panel the amount of S segment RNA released from PIwUHV cells. **C)** The left panel shows the amount of L and the right panel the amount of S segment RNA released from PIwUGV-1 cells. The black arrows on the right-hand side indicate the amount of persistently infecting virus released (S segment RNA) during eight days incubation in the absence of superinfection.

## DISCUSSION

Boa constrictors with BIBD most often carry multiple reptarenavirus S and L segments (26, 27) that are genetically distant from each other. Similar coinfections are not common with mammarenaviruses, although a recent report described the presence of two genetically divergent L segments with a single S segment in a wild mouse (33). In line with the former, studies on persistently mammarenavirus infected cell cultures have demonstrated resistance against superinfection with closely related mammarenaviruses (34–36). These mammarenavirus studies and our observations from natural reptarenavirus coinfections led us to hypothesize that reptarenavirus S and L segments classified into the same species would compete in replication during coinfection, while reptarenaviruses of different species would not. To test the hypothesis, we performed a set of coinfection experiments on kidney- and brain derived boa constrictor cell lines (I/1Ki and V/4Br) with one hartmanivirus and five reptarenavirus isolates. In addition, we tested the ability of the virus isolates to superinfect cell cultures persistently infected with one or two reptarenaviruses, or with reptarenavirus and hartmanivirus.

We started off by comparing the amount of L and S segment released into the supernatant in coinfections versus single-virus infections in the boa constrictor kidney (I/1Ki) brain (V/4Br) cell lines. Overall, coinfection did not appear to markedly affect the amount of both L and S segments released from the I/1Ki cells, which can efficiently be infected with all of the virus isolates included. However, coinfection of I/1Ki cells with UHV-1 and UHV-2, which by sequence homology would belong to the same reptarenavirus species, showed a decrease in the amount of released UHV-1 segments. The brain-derived V/4Br cells appeared to be less potent in releasing viral RNA of the selected viruses. The observations could be explained by e.g. the cells being less permissive or less capable of maintaining replication. The results are in line with the findings of our earlier study which indicated that I/1Ki cells are more permissive towards reporter-bearing vesicular stomatitis viruses pseudotyped with reptarenavirus glycoproteins than V/4Br cells (32). A less efficient entry could explain the observed 100-to 10,000-fold difference in the amount of viral RNA released from the V/4Br cells. In terms of coinfection versus single virus infection, the results suggested, like in I/1Ki cells, that coinfection did not markedly affect the amount of viral RNA released for the majority of viruses. However, as seen in I/1Ki cells, the release of UHV-1 segments was reduced in coinfections with UHV-2. Interestingly, the release of UHV-1 segments also appeared to be slightly affected by coinfection with other reptarenaviruses in this cell line.

Because the results with UHV-1 and UHV-2 coinfections appeared to align with our initial hypothesis of competition between closely related viruses, we decided to make use of the persistently infected (PI) cell cultures of our earlier study (18). We first showed that the persistently infected cell lines can be superinfected with all tested viruses, and that superinfection of the PI cells did not markedly affect the amount of RNA released of the persistently infecting virus in most virus combinations, although most cell lines showed a slight increase (up to approximately 2-fold) in the amount of released RNA of the persistently infecting virus. The enhanced release was most obvious for UGV-1, which showed an over 10-fold increase in the amount of released RNA. The enhancement of the RNA release following superinfection merits further studies to properly document the putative enhancement and reveal its mechanism. When establishing the PI cultures, we observed a gradual decline in the S segment encoded proteins NP and the glycoproteins over the first 10 passages (18). This could indicate that during persistent infection the fitness of the virus declines or that for some reason the expression level of the viral segments is lower. The increased RNA release following superinfection could thus indicate a “helper” function of the superinfecting virus, i.e. that the RNA of the persistently infecting virus would be released also in particles bearing the GPs of the superinfecting virus. A comparison of superinfections with co- and monoinfections in naïve I/1Ki cells showed that similar amounts of viral RNA were released in most virus combinations. However, superinfection with a closely related virus, i.e. UGV-2 in the case of PIwUGV-1 (persistently infected with UGV-1), UHV-1 in the case of PIwSn11 (persistently infected with UHV-2 and HISV-1) and UHV-2 in the case of PIwUHV (persistently infected with ABV-1 and UHV-1), resulted in a clearly attenuated release of the superinfecting virus’ RNA. These results are well in line with the initial hypothesis of competition between closely related but not with more distantly related reptarenaviruses. They would thus help to explain the observation that snakes with BIBD do not carry several segments of the same reptarenavirus species. Furthermore, they align well with the cell culture studies on mammarenaviruses which have shown resistance of persistently infected cultures towards superinfection with closely related mammarenaviruses (34–36). The mechanism of the putative superinfection exclusion of closely related viruses requires further studies, however, we propose that RdRp-ZP interaction would be the mediating factor. In mammarenaviruses, RdRp and ZP form a complex in a species-specific manner and the complex formation regulates the RNA synthesis (37). Such a mechanism would directly fit the observations of superinfection exclusion between viruses of the same species. RdRp-ZP interaction could also explain the coinfection competition between viruses of the same species; the virus that produces ZP more efficiently (or a virus whose ZP is more suitable for interaction between both RdRps) would eventually win the competition.

While establishing the persistently reptarenavirus infected cell lines, we observed rather large differences in the ratio of S and L segments released depending on the virus isolate (18), which motivated us to look at the ratio of released S/L segment RNA also in the present study. In line with the previous study, we found that the cells infected with different virus isolates release varying S and L segment ratios. With all tested viruses, except UHV-2 (ratio close to 1), the S/L segment ratio remained above 2 in I/1Ki cells, indicating that the cells release at least two S segments per each L segment. For most viruses tested the V/4Br cells showed similar ratios, suggesting that the ratio of released S/L segment might not be cell type dependent. Instead, we think that the differences in the ratio of segments released from I/1Ki versus V/4Br cells observed for some viruses would rather relate to the inability of these viruses to establish a productive infection in the V/4Br cells.

Interestingly, some virus combinations in both superinfection and coinfection, e.g. UHV-1 with UGV-1 or UGV-2 (also UGV-1 with UHV-1) and UGV-2 with UHV-2 or UGV-1, showed an altered ratio of the released segments. Furthermore, the ratio appeared to vary depending on the sampling time point, indicating that the dynamics between and the levels of the segments might fluctuate during infection. While the results do not provide information on the number of S and L segments within a single particle, the rather large variability in the size of arenavirus particles (8) could allow the release of e.g. two S segments and one L segment or even two L segments and one S segment in a single particle. In the case of coinfection this could theoretically allow the release of particles with genetically divergent S or L segments. Further studies on the virion’s RNA composition would have merits since it might have implications on the spread of reptarenaviruses and the pathogenesis of BIBD. All things considered, the results of the present study provide cell culture evidence to support the hypothesis that consecutive reptarenavirus superinfections among captive snake populations has led to the accumulation of genetically divergent reptarenavirus S and L segments.

## ACKNOWLEDGEMENTS

The authors wish to acknowledge Dr. Antti Hassinen from FIMM (Institute for Molecular Medicine Finland High Content Imaging and Analysis unit) for his help with Opera Phenix High Content Screening system. The study was supported by the Academy of Finland (J.H.; grants 308613, 314119, and 335762). The funder had no role in designing the study, or in interpretation and presentation of the results.

## REFERENCES

1. Rohwer F, Barott K. 2013. Viral information. Biology & Philosophy 28:283–297.

2. Kumar N, Sharma S, Barua S, Tripathi Bhupendra N, Rouse Barry T. 2018. Virological and Immunological Outcomes of Coinfections. Clinical Microbiology Reviews 31:e00111–17.

3. Griffiths EC, Pedersen AB, Fenton A, Petchey OL. 2011. The nature and consequences of coinfection in humans. J Infect 63:200–6.

4. Abudurexiti A, Adkins S, Alioto D, Alkhovsky SV, Avsic-Zupanc T, Ballinger MJ, Bente DA, Beer M, Bergeron E, Blair CD, Briese T, Buchmeier MJ, Burt FJ, Calisher CH, Chang C, Charrel RN, Choi IR, Clegg JCS, de la Torre JC, de Lamballerie X, Deng F, Di Serio F, Digiaro M, Drebot MA, Duan X, Ebihara H, Elbeaino T, Ergunay K, Fulhorst CF, Garrison AR, Gao GF, Gonzalez JJ, Groschup MH, Gunther S, Haenni AL, Hall RA, Hepojoki J, Hewson R, Hu Z, Hughes HR, Jonson MG, Junglen S, Klempa B, Klingstrom J, Kou C, Laenen L, Lambert AJ, Langevin SA, Liu D, Lukashevich IS, et al. 2019. Taxonomy of the order Bunyavirales: update 2019. Archives of Virology 164:1949.

5. Salazar-Bravo J, Ruedas LA, Yates TL. 2002. Mammalian Reservoirs of Arenaviruses, p 25–63. In Oldstone Mba (ed), Arenaviruses I: The Epidemiology, Molecular and Cell Biology of Arenaviruses doi:10.1007/978-3-642-56029-3_2. Springer Berlin Heidelberg, Berlin, Heidelberg.

6. Downs WG, Anderson CR, Spence L, Aitken TH, Greenhall AH. 1963. Tacaribe virus, a new agent isolated from Artibeus bats and mosquitoes in Trinidad, West Indies. The American Journal of Tropical Medicine and Hygiene 12:640.

7. Sayler KA, Barbet AF, Chamberlain C, Clapp WL, Alleman R, Loeb JC, Lednicky JA. 2014. Isolation of Tacaribe virus, a Caribbean arenavirus, from host-seeking Amblyomma americanum ticks in Florida. PLoS One 9:e115769.

8. Radoshitzky SR, Buchmeier MJ, Charrel RN, Clegg JCS, Gonzalez JJ, Gunther S, Hepojoki J, Kuhn JH, Lukashevich IS, Romanowski V, Salvato MS, Sironi M, Stenglein MD, de la Torre JC, Ictv Report C. 2019. ICTV Virus Taxonomy Profile: Arenaviridae. The Journal of general virology 100:1200.

9. Hepojoki J, Hepojoki S, Smura T, Szirovicza L, Dervas E, Prahauser B, Nufer L, Schraner EM, Vapalahti O, Kipar A, Hetzel U. 2018. Characterization of Haartman Institute snake virus-1 (HISV-1) and HISV-like viruses-The representatives of genus Hartmanivirus, family Arenaviridae. PLoS pathogens 14:e1007415.

10. Radoshitzky SR, Bao Y, Buchmeier MJ, Charrel RN, Clawson AN, Clegg CS, DeRisi JL, Emonet S, Gonzalez JP, Kuhn JH, Lukashevich IS, Peters CJ, Romanowski V, Salvato MS, Stenglein MD, de la Torre JC. 2015. Past, present, and future of arenavirus taxonomy. Archives of Virology 160:1851.

11. Hallam SJ, Koma T, Maruyama J, Paessler S. 2018. Review of Mammarenavirus Biology and Replication. Front Microbiol 9:1751.

12. Stenglein MD, Sanchez-Migallon Guzman D, Garcia VE, Layton ML, Hoon-Hanks LL, Boback SM, Keel MK, Drazenovich T, Hawkins MG, DeRisi JL. 2017. Differential Disease Susceptibilities in Experimentally Reptarenavirus-Infected Boa Constrictors and Ball Pythons. Journal of virology 91:10.1128/JVI.00451.

13. Stenglein MD, Sanders C, Kistler AL, Ruby JG, Franco JY, Reavill DR, Dunker F, Derisi JL. 2012. Identification, characterization, and in vitro culture of highly divergent arenaviruses from boa constrictors and annulated tree boas: candidate etiological agents for snake inclusion body disease. mBio 3:e00180.

14. Bodewes R, Kik MJ, Raj VS, Schapendonk CM, Haagmans BL, Smits SL, Osterhaus AD. 2013. Detection of novel divergent arenaviruses in boid snakes with inclusion body disease in The Netherlands. The Journal of general virology 94:1206.

15. Hetzel U, Korzyukov Y, Keller S, Szirovicza L, Pesch T, Vapalahti O, Kipar A, Hepojoki J. 2021. Experimental Reptarenavirus Infection of Boa constrictor and Python regius. Journal of virology 95:e01968.

16. Hetzel U, Sironen T, Laurinmäki P, Liljeroos L, Patjas A, Henttonen H, Vaheri A, Artelt A, Kipar A, Butcher SJ, Vapalahti O, Hepojoki J. 2013. Isolation, Identification, and Characterization of Novel Arenaviruses, the Etiological Agents of Boid Inclusion Body Disease. Journal of Virology 87:10918--10935.

17. Schumacher J, Jacobson ER, Homer BL, Gaskin JM. 1994. Inclusion Body Disease in Boid Snakes. Journal of Zoo and Wildlife Medicine 25:511.

18. Lintala A, Szirovicza L, Kipar A, Hetzel U, Hepojoki J. 2022. Persistent Reptarenavirus and Hartmanivirus Infection in Cultured Boid Cells. Microbiol Spectr doi:10.1128/spectrum.01585-22:e0158522.

19. Keller S, Hetzel U, Sironen T, Korzyukov Y, Vapalahti O, Kipar A, Hepojoki J. 2017. Coinfecting Reptarenaviruses Can Be Vertically Transmitted in Boa Constrictor. PLoS Pathogens 13.

20. Windbichler K, Michalopoulou E, Palamides P, Pesch T, Jelinek C, Vapalahti O, Kipar A, Hetzel U, Hepojoki J. 2019. Antibody response in snakes with boid inclusion body disease. PloS one 14:e0221863.

21. Chang L, Fu D, Stenglein MD, Hernandez JA, DeRisi JL, Jacobson ER. 2016. Detection and prevalence of boid inclusion body disease in collections of boas and pythons using immunological assays. Veterinary journal (London, England : 1997) 218:13.

22. Aqrawi T, Stohr AC, Knauf-Witzens T, Krengel A, Heckers KO, Marschang RE. 2015. Identification of snake arenaviruses in live boas and pythons in a zoo in Germany. Tierarztliche PraxisAusgabe K, Kleintiere/Heimtiere 43:239.

23. Dietz J, Kolesnik E, Heckers KO, Marc-Niklas K, Marschang RE. 2020. Detection of an Arenavirus in a Group of Captive Wagler’s Pit Vipers (Tropidolaemus Wagleri). Journal of Zoo and Wildlife Medicine 51:236.

24. Hyndman TH, Marschang RE, Bruce M, Clark P, Vitali SD. 2019. Reptarenaviruses in apparently healthy snakes in an Australian zoological collection. Australian Veterinary Journal 97:93.

25. Simard J, Marschang RE, Leineweber C, Hellebuyck T. 2020. Prevalence of inclusion body disease and associated comorbidity in captive collections of boid and pythonid snakes in Belgium. PloS one 15:e0229667.

26. Hepojoki J, Salmenpera P, Sironen T, Hetzel U, Korzyukov Y, Kipar A, Vapalahti O. 2015. Arenavirus Coinfections Are Common in Snakes with Boid Inclusion Body Disease. Journal of Virology 89:8657--8660.

27. Stenglein MD, Jacobson ER, Chang LW, Sanders C, Hawkins MG, Guzman DS, Drazenovich T, Dunker F, Kamaka EK, Fisher D, Reavill DR, Meola LF, Levens G, DeRisi JL. 2015. Widespread recombination, reassortment, and transmission of unbalanced compound viral genotypes in natural arenavirus infections. PLoS pathogens 11:e1004900.

28. Argenta FF, Hepojoki J, Smura T, Szirovicza L, Hammerschmitt ME, Driemeier D, Kipar A, Hetzel U. 2020. Identification of Reptarenaviruses, Hartmaniviruses and a Novel Chuvirus in Captive Brazilian Native Boa Constrictors with Boid Inclusion Body Disease. Journal of virology doi:JVI.00001-20 [pii].

29. Szirovicza L, Hetzel U, Kipar A, Martinez-Sobrido L, Vapalahti O, Hepojoki J. 2020. Snake Deltavirus Utilizes Envelope Proteins of Different Viruses To Generate Infectious Particles. mBio 11:10.1128/mBio.03250.

30. Alfaro-Alarcón A, Hetzel U, Smura T, Baggio F, Morales JA, Kipar A, Hepojoki J. 2022. Boid Inclusion Body Disease Is Also a Disease of Wild Boa Constrictors. Microbiol Spectr doi:10.1128/spectrum.01705-22:e0170522.

31. Chang L-W, Jacobson ER. 2010. Inclusion Body Disease, A Worldwide Infectious Disease of Boid Snakes: A Review. Journal of Exotic Pet Medicine 19:216.

32. Korzyukov Y, Iheozor-Ejiofor R, Levanov L, Smura T, Hetzel U, Szirovicza L, de la Torre JC, Martinez-Sobrido L, Kipar A, Vapalahti O, Hepojoki J. 2020. Differences in Tissue and Species Tropism of Reptarenavirus Species Studied by Vesicular Stomatitis Virus Pseudotypes. Viruses 12:10.3390/v12040395.

33. Cuypers LN, Čížková D, de Bellocq JG. 2022. Co-infection of mammarenaviruses in a wild mouse, Tanzania. Virus Evolution 8:veac065.

34. Ellenberg P, Edreira M, Scolaro L. 2004. Resistance to superinfection of Vero cells persistently infected with Junin virus. Archives of Virology 149:507.

35. Ellenberg P, Linero FN, Scolaro LA. 2007. Superinfection exclusion in BHK-21 cells persistently infected with Junin virus. The Journal of general virology 88:2730.

36. Damonte EB, Mersich SE, Coto CE. 1983. Response of cells persistently infected with arenaviruses to superinfection with homotypic and heterotypic viruses. Virology 129:474.

37. Kranzusch PJ, Whelan SP. 2011. Arenavirus Z protein controls viral RNA synthesis by locking a polymerase-promoter complex. Proceedings of the National Academy of Sciences of the United States of America 108:19743.

